# Isolation and comparative genomic analysis of reuterin-producing *Lactobacillus reuteri* from poultry gastrointestinal tract

**DOI:** 10.1101/793299

**Authors:** Anna Greppi, Paul Tetteh Asare, Clarissa Schwab, Niklaus Zemp, Roger Stephan, Christophe Lacroix

**Author notes:** The authors equally contributed to the manuscript (Anna Greppi, Paul Tetteh Asare).

## Abstract

*Lactobacillus reuteri* is a natural inhabitant of selected animal and human gastrointestinal tract (GIT). Certain strains have the capacity to transform glycerol to 3-hydroxypropionaldehyde (3-HPA), further excreted to form reuterin, a potent antimicrobial system. Reuterin-producing strains may be applied as a natural antimicrobial in feed to prevent pathogen colonization of animals, such as in poultry, and replace added antimicrobials. To date, only seven *L. reuteri* strains isolated from poultry have been characterized which limits phylogenetic studies and host-microbes interactions characterization. This study aimed to isolate *L. reuteri* strains from poultry GIT and to characterize their reuterin production and antimicrobial resistance (AMR) profiles using phenotypic and genetic methods. Seventy reuterin-producing strains were isolated from poultry crop, faeces and caeca and twenty-five selected for further characterization. Draft genomes were generated for the new 25 isolates and integrated in a phylogenetic tree of 40 strains from different hosts. Phylogenetic analysis based on gene content as well as on core genomes showed grouping of the selected 25 *L. reuteri* poultry isolates within the poultry/human lineage VI. Strains harbouring *pdu-cob-cbi-hem* genes (23/25) produced between 156 mM ± 11 and 330 mM ± 14 3-HPA, from 600 mM of glycerol, in the conditions of the test. All 25 poultry strains were sensitive to cefotaxime (MIC between 0.016 and 1 μg/mL) and penicillin (MIC between 0.02 and 4 μg/mL). Akin to the reference strains DSM20016 and SD2112, the novel isolates were resistant to penicillin, possibly associated with identified point mutations in *ponA*, *pbpX*, *pbpF* and *pbpB*. All strains resistant to erythromycin (4/27) carried the ermB gene, and it was only present in poultry strains. All strains resistant to tetracycline (5/27) harbored tetW gene. This study confirms the evolutionary history of poultry/human lineage VI and identifies *pdu-cob-cbi-hem* as a frequent trait but not always present in this lineage. *L. reuteri* poultry strains producing high 3-HPA yield may have potential to prevent enteropathogen colonization of poultry.

## Introduction

*Lactobacillus reuteri* inhabits the gastrointestinal tract (GIT) of selected animals where it forms biofilms on the non-glandular, squamous epithelium lining the upper GIT. In poultry, *L. reuteri* is the most abundant *Lactobacillus* species in the GIT, mainly found in the crop and the caecum [1]. Distinct phylogenetic lineages of *L. reuteri* are coherent with host origin, reflecting co-evolution of this species with the vertebrate hosts [2]. The evolutionary adaptation differentiates the species in host-adapted phylogenetic lineages comprised of isolates from rodents (lineages I and III), humans (lineage II), pigs (lineages IV and V) and poultry/human (lineage VI) [2, 3]. Host adaption has been linked to the occurrence of specific functional traits, e.g. rodent *L. reuteri* isolates possess the genes responsible for synthesis of urease, as the strains are constantly exposed to urea in the forestomach of mice [4].

Genomes of poultry and human *L. reuteri* isolates (lineages II and VI) have been shown to harbour the *pdu-cbi-cob-hem* operon, as a lineage specific trait [5]. This operon contains genes for glycerol and propanediol utilization (*pdu*) and for cobalamin biosynthesis (*cbi-cob*), *hem* genes and some accessory genes. Cobalamin is a co-factor for glycerol/diol dehydratase PduCDE (EC 4.2.1.30). PduCDE catalyzes the conversion of 1,2-propanediol to propanal, which can be further metabolized by other enzymes of the *pdu* operon to propanol or propionate [6]. Glycerol, a second substrate of PduCDE, is transformed to the intermediate 3-hydroxypropionaldehyde (3-HPA) which can be further metabolized to 1,3-propanediol or 3-hydroxypropionate [7]. 3-HPA produced from glycerol is released from the cell forming the dynamic multi-compound reuterin system, with broad antimicrobial spectrum and consisting of 3-HPA, its hydrate and dimer and acrolein [8, 9]. Acrolein, a highly reactive toxicant, was recently shown to be the main component for the antimicrobial activity of reuterin [10, 11].

Due to the high persistence of *L. reuteri* in the poultry GIT and the established antimicrobial activity of reuterin, *L. reuteri* has high potential to be applied as a natural antimicrobial in feed to prevent pathogen infection of animals [12]. *L. reuteri* strains isolated from poultry GIT were shown effective against *Salmonella* spp. and *Escherichia coli* resistant to various antibiotics [13]. Moreover, *L. reuteri* in the early post-hatching period had a delayed effect on ileum microbiota of poultry, which resulted in the enrichment of potentially beneficial lactobacilli and the suppression of Proteobacteria [14]. To select functional *L. reuteri* strains, a key trait is the determination of their antimicrobial resistance (AMR) profiles to identify intrinsic and extrinsic resistances that may be potentially transferred. Lactobacilli are known to be intrinsically resistant against vancomycin. However, the occurrence of tetracycline and erythromycin genes on mobile elements has been reported for different *Lactobacillus* spp. [15].

To date, among all deposited NCBI *L. reuteri* genomes, only seven strains were isolated from poultry, representing only 5 % of the isolates and thus limiting any phylogenetic analysis and host-microbes adaptation studies. The majority of the NCBI *L. reuteri* deposited genomes comes from strains which had been isolated from mouse (47), human (19) and pig (25), while few from sourdough (7), poultry (7), goat (5), cow (5), rat (4) sheep (4), dairy and fermented products (3), horse (3), piglet (2), pork (1), probiotic capsule (1), wine (1) and yoghurt (1).

It was therefore the aim of this study to isolate and characterize *L. reuteri* strains from poultry and characterize their reuterin production and AMR profiles using phenotypic and genotypic methods. Draft genomes of the isolates were analyzed combined with 40 *L. reuteri* genomes of strains previously isolated from different hosts to assess genetic diversity and gain insight into distinguishing features related to poultry, and enrich previously phylogenetic characterization of *L. reuteri*.

## Material and methods

### Bacterial strains and growth conditions

*L. reuteri* DSM20016 (Leibniz Institute DSMZ-German Collection of Microorganisms and Cell Cultures, Braunschweig, Germany) and *L. reuteri* SD2112 (BioGaia AB, Stockholm, Sweden) were used as reference strains. Reference strains as well as all *L. reuteri* isolated in this study were propagated anaerobically (Oxoid, AnaeroGenTM, Basingstoke, UK) at 37°C in de Man, Rogosa and Sharpe (MRS) broth medium (Biolife, Milan, Italy).

### Bacterial isolation

Six Lohmann brown poultry (13 weeks old) were obtained from six poultry farms in Switzerland. Crop, caeca and faeces were aseptically collected. In parallel, 10 whole gut of Cobb 500 broiler poultry were obtained from Schönholzer Werner abattoir in Wädenswil (Zurich), and transported to the lab within 1 hour.

*L. reuteri* strains were isolated from poultry crop using the protocol previously described [4], with some modifications. Briefly, one gram of crop, caecal or faecal content was added to 10 mL of sterile phosphate buffer saline (PBS) (137 mM NaCl, 2.7 mM KCl, 10 mM Na_2_HPO_4_, 2 mM KH_2_PO_4_, pH 7.4) and homogenized in a stomacher (BagMixer® 400 P, Interscience, Saint Nom, France) at high speed for 1 min. Suspensions from samples were serially diluted and spread on mMRS plates [16]. The agar plates were incubated overnight at 42 °C under anaerobic condition using AnaeroGen 2.5 L (Thermo Fisher Diagnostics AG, Pratteln, Switzerland). Replica plates were prepared using Scienceware replica plater and velveteen squares (Sigma-Aldrich, Buchs, Switzerland), and incubated overnight as presented above. After incubation, one plate was overlaid with 500 mM glycerol agar (1% agar) and incubated at 37 °C for 30 min for testing reuterin production of colonies. A colorimetric method was used with the addition of 5 mL 2,4-dinitrophenylhydrazine (0.1% in 2 M HCl), 3 min incubation, removal of the solution, addition of 5 mL 5M KOH. *L. reuteri* colonies showing purple zones indicating reuterin synthesis were streaked on MRS agar plates and single colony were sub cultured 3 times in MRS broth (1% inoculum, 18 h at 42 °C). Few negative colonies which show a colony morphology of *L. reuteri* but no purple zone were picked as negative controls. Species confirmation and reuterin production quantification was performed for both positive and selected negative selected colonies.

### Bacterial identification

Genomic DNA was isolated using a lysozyme-based cell wall digestion followed by the Wizard genomic DNA purification kit (Promega, Dübendorf, Switzerland). Total DNA was quantified by absorbance at 260 nm using NanoDrop® ND-1000 Spectrophotometer (Witec AG, Littau, Switzerland). DNA quality was analyzed by electrophoresis in 1.2% (w/v) agarose gel and Gel Red staining (VWR International AG, Dietikon, Switzerland). The DNA samples were stored at −20 °C until further analysis.

To confirm the identity of the isolates, 1.6 kbp full region of 16S rRNA gene was amplified by PCR using universal primers bak4 (5’-AGGAGGTGATCCARCCGCA-3’) and bak11w (5’-AGTTTGATCMTGGCTCAG-3’) [17, 18]. The 16S rRNA PCR assay consisted of 5 min at 95 °C, followed by 35 cycles of 15 s at 95 °C, 30 s at 60 °C and 2 min at 72 °C and final extension for 7 min at 72 °C. Sanger sequencing of the PCR amplicon was performed at GATC (Konstanz, Germany). To identify the closest homologs, DNA sequences obtained were aligned using Basic Local Alignment Search Tool (BLAST) [19]. Sequence homology greater than 97% was used to identify *L. reuteri*.

### Strain typing

The enterobacterial repetitive intergenic consensus (ERIC) sequence was used to differentiate between *L. reuteri* isolates, as previously described and using ERIC1R (5’ ATGTAAGCTCCTGGGGATTCAC-3’) and ERIC2 (5’ AAGTAAGTGACTGGGGTGAGCG-3’) primers [20–22]. The ERIC-PCR assay was performed using 100 ng of template DNA of each isolate. The protocol consisted of 7 min at 95 °C, followed by 30 cycles of 30 s at 90 °C, 1 min at 52 °C, and 8 min at 65 °C, and a final extension for 16 min at 65 °C. The ERIC-PCR amplicons were analyzed on 2% (wt/vol) agarose gels for 6 h at 60 Volts. GeneRuler DNA ladder mix (Fermentas, Le Mont-sur-Lausanne, Switzerland) was used as a molecular size marker according to the manufacturer’s directions and gels were visualized by Gel Red staining. Gels were analyzed using Gel Compar II version 6.5 software package (Applied Maths, Sint-Martens-Latem, Belgium). *L. reuteri* strains with unique ERIC profiles were visually selected and amplicons re-run on a single agarose gel. A dendrogram of similarity was generated from the gel with selected isolates using the Pearson correlation similarity coefficient and the unweighted-pair group method (UPGMA) with arithmetic averages, and 1% optimization. Based on the clustering obtained, confirmed isolates with unique profiles were selected for whole genome sequencing and characterization.

### Generation and annotation of draft genomes

Genomic DNA from the isolated strains was obtained as described above and standardized to 100 ng/μL. The whole genomes of 25 *L. reuteri* isolates were sequenced with the standard set of 96 Illumina paired-end barcodes on a HiSeq 2500 Illumina Technology (Illumina Inc., San Diego, USA) with 2× 125 high output mode. The genomic library was generated using reagents from NEBNext Illumina preparation kit. Raw paired-end reads were quality trimmed using the default settings of Trimmomatic [23]. Read pairs were merged by FLASH [24]. De novo assembly was performed with SPAdes assembler (version 3.12) with –*careful* option [25]. The quality of the assembly based on evolutionarily-informed expectations of gene content from near-universal single-copy orthologs selected from OrthoDB v9 was assessed by BUSCO [26] and QUAST [27]. Reference guided ordering of scaffolds based on iterative alignment steps was performed by QUAST using *L. reuteri* DSM20016 genome as reference. SeqKit [28] and QUAST were used to retrieve genome features. The complete genome of *L. reuteri* DSM20116 type strain was compared individually with each of the 25 draft genomes of *L. reuteri* poultry isolates. Average nucleotide identity (ANI) values between two genomic datasets were calculated using JSpecies [29]. Genomes with ANI values above 95 % were considered as belonging to the same species [30].

Twenty-five *L. reuteri* draft genomes (this study) and 40 NCBI deposited *L. reuteri* genomes of strains isolated from different hosts (poultry, human, mouse, rat, pig, sourdough, goat, sheep, cow, horse; Supplementary Table S1) were structurally annotated using the PROKKA 3.10.1 suite [31]. Only contigs higher than 500 bp were included in the analysis, and *Lactobacillus* was selected as reference database for the annotation (options used: --*genus* Lactobacillus --*species* reuteri --*usegenus* Lactobacillus –*mincontiglen* 500).

### Comparative genomics

Comparative genome analysis was based on gene content tree, core genome phylogenetic tree and a nucleotide-content similarity matrix (ANI matrix). For the generation of the gene content tree, a matrix based on gene content (binary data for presence or absence of each annotated gene) was generated comprising all annotated genes of 25 *L. reuteri* draft genomes plus the 40 *L. reuteri* NCBI genomes. The gene content tree was then constructed using the hierarchical cluster analysis (hcust) on R 3.4.4 while a core genome phylogenetic tree was calculated using EDGAR 2.3 [32]. Concatenated sequences were used to calculate a distance matrix which provided the input for the neighbour-joining method with PHYLIP implementation. EDGAR 2.3 calculated the core genome of each identified clusters as a set of orthologous genes present in all strains belonging to each cluster. For the generation of the similarity matrix, ANI was calculated on EDGAR 2.3 with an all-against-all comparisons at nucleotide level for all 65 *L. reuteri* strains.

Identified lineages were named based on host origin of the strains, and lineages specific features were determined using “Define metacontigs” function in EDGAR 2.3. Core groups were created for the sets of strains derived from the phylogenetic analysis. A Venn diagram was then designed in EDGAR to identify shared and unique genes among the group of strains. Unique genes of poultry/human lineage VI were manually categorized based on UniProt protein description into the following groups: transport proteins, DNA-binding proteins, transferases, lyases, oxidoreductases, membrane proteins, hydrolases, virulence related proteins, RNA-binding and prophage-related proteins. The presence of genes of the *pdu-cob-hem-cbi* cluster was manually checked and a heatmap of presence/absence of genes of interest was designed with GraphPad Prism 8.0 (GraphPad Software Inc. La Jolla California, USA). The presence of AMR genes in the assembled genomes was also manually checked and integrated with the results from ResFinder 3.0 tool [33].

For penicillin resistant *L. reuteri* strains, point mutations (SNPs) in penicillin-binding proteins genes *ponA, pbpX, pbpF* and *pbpB* were checked by aligning deduced amino acid sequences of resistant and sensitive strains with NCBI-deposited sequences for *Lactobacillus* in Geneious 9.1.8.

### PCR for *tetW* and *ermB* detection

PCR for *tetW* and *ermB* genes was performed to confirm the genomic data. *TetW* was amplified with tetw-rev and tetw-fw primers [15, 34]. The PCR protocol comprised 35 cycles of 95 °C for 30 s, 56 °C 45 and 72 °C 30 s. The *ermB* gene was amplified with ermA1 and ermA2 primers using 30 PCR cycles composed of 95 °C for 30 s, 52 °C 45 s, 72 °C 2 min [15].

### Reuterin production

Reuterin production was determined using a two-step process [35]. Cell pellets obtained from 16 h *L. reuteri* cultures (OD_600_ approx. 8.0) were collected by centrifugation, resuspended in 600 mM glycerol solution and incubated at 25 °C for 2 h. For reuterin biosynthesis, the concentrations of glycerol, 3-HPA and acrolein were measured by high performance liquid chromatography using refractive index (HLPC-RI, Hitachi LaChrome, Merck, Dietikon, Switzerland) and ion-exclusion chromatography with pulsed-amperometric detection (IC-PAD) analysis, respectively [8, 11]. For HPLC-RI, an Aminex HPx-87H column and sulfuric acid (10 mM) was used as eluent. The IC-PAD Thermo Scentific (Reinach, Switzerland) ICS-5000+ system was equipped with a quaternary gradient pump, a thermostated autosampler and an electrochemical detector with a cell containing a Ag/AgCl reference electrode and a disposable thin film platinum working electrode tempered at 25°C. Analytes were separated with a Thermo Scientific IonPac ICE-AS1 4 × 250 mm ion-exclusion column with a guard column, operated at 30°C. The solvent system was isocratic 0.1 M methanesulfonic acid at 0.2 mL/min for 36 min.

### Antimicrobial sensitivity profiling

*L. reuteri* poultry isolates and reference strains *L. reuteri* SD2112 and DSM20016 were tested for susceptibility to cefotaxime (CFX), erythromycin (ERM), penicillin (PEN), tetracycline (TET), vancomycin (VAN) and ciprofloxacin (CIP), using gradient diffusion MTS™ strips (Liofilchem, Roseto degli Abruzzi, Italy). The concentration range tested was 0.016 to 256 μg/mg, except for CIP (0.002 to 32 μg/mL). Briefly, 16 h overnight *L. reuteri* cultures were added to a sterile MRS broth (1% v/v) and incubated at 37 °C for 6 h. The bacterial cultures were standardized to OD_600_ 1.0 (corresponding to approximately 10^8^ CFU/mL) using a PowerWave XS microplate spectrophotometer (BioTek, Sursee, Switzerland). The diluted culture (10^6^ CFU/mL) was evenly swabbed on MRS agar plates in duplicate using sterile cotton bud. MTS™ strips were placed on the surface of the agar and incubated at 37 °C for 24 h in anaerobic condition supplied by gas package (AnaeroGen, Thermo Fisher Diagnostics AG, Pratteln, Switzerland). The minimum inhibitory concentration (MIC) were recorded as the point where inhibition curves intersect the scale on the MTS™ strip.

### Data Accession Number

The draft genomes of 25 *L. reuteri* strains have been deposited on NCBI under the accession numbers indicated in Table 1. Genome sequences are available in the GenBank under the BioProject ID: PRJNA473635.

**Table 1.**
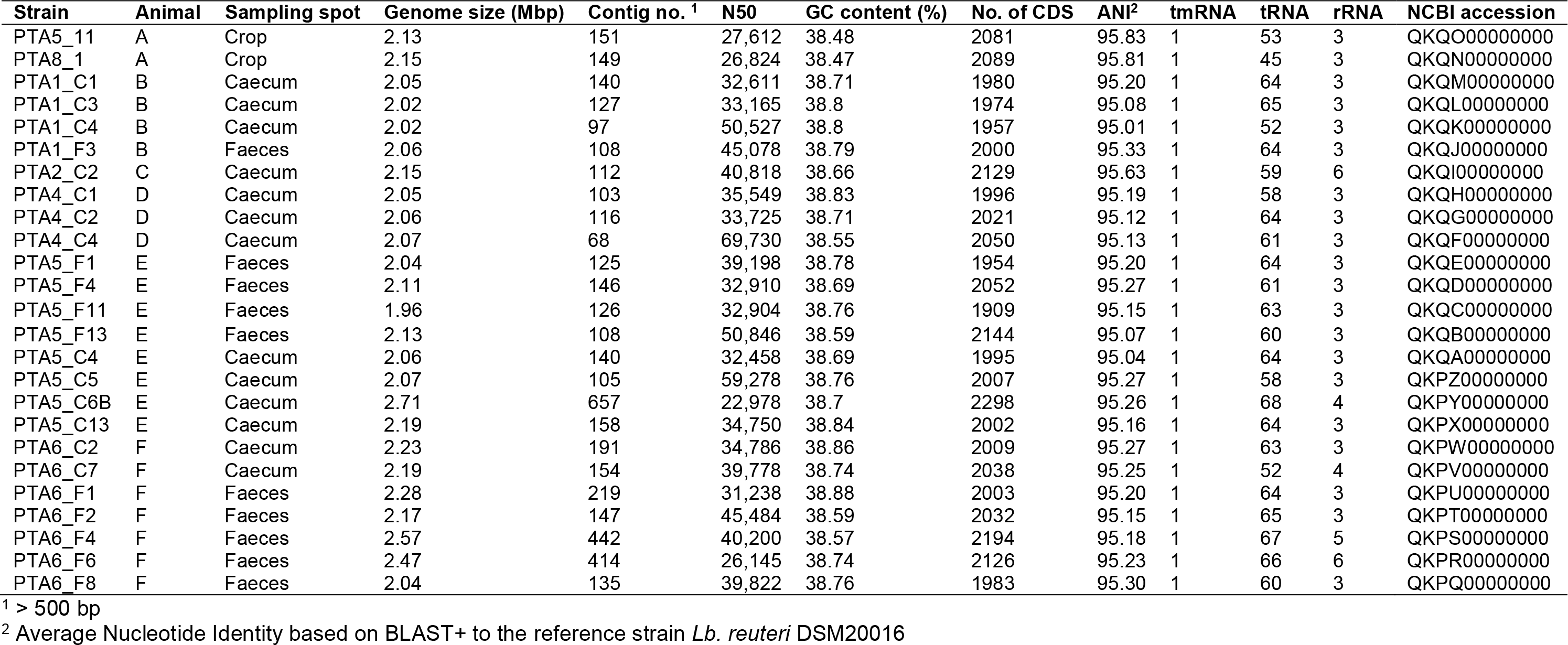
Draft genome features of 25 *L. reuteri* strains isolated from poultry GIT and sequenced in this study.

| Strain | Animal | Sampling spot | Genome size (Mbp) | Contig no. <sup>1</sup> | N50 | GC content (%) | No. of CDS | ANI<sup>2</sup> | tmRNA | tRNA | rRNA | NCBI accession |
| --- | --- | --- | --- | --- | --- | --- | --- | --- | --- | --- | --- | --- |
| PTA5_11 | A | Crop | 2.13 | 151 | 27,612 | 38.48 | 2081 | 95.83 | 1 | 53 | 3 | QKQO00000000 |
| PTA8_1 | A | Crop | 2.15 | 149 | 26,824 | 38.47 | 2089 | 95.81 | 1 | 45 | 3 | QKQN00000000 |
| PTA1_C1 | B | Caecum | 2.05 | 140 | 32,611 | 38.71 | 1980 | 95.20 | 1 | 64 | 3 | QKQM00000000 |
| PTA1_C3 | B | Caecum | 2.02 | 127 | 33,165 | 38.8 | 1974 | 95.08 | 1 | 65 | 3 | QKQL00000000 |
| PTA1_C4 | B | Caecum | 2.02 | 97 | 50,527 | 38.8 | 1957 | 95.01 | 1 | 52 | 3 | QKQK00000000 |
| PTA1_F3 | B | Faeces | 2.06 | 108 | 45,078 | 38.79 | 2000 | 95.33 | 1 | 64 | 3 | QKQJ00000000 |
| PTA2_C2 | C | Caecum | 2.15 | 112 | 40,818 | 38.66 | 2129 | 95.63 | 1 | 59 | 6 | QKQI00000000 |
| PTA4_C1 | D | Caecum | 2.05 | 103 | 35,549 | 38.83 | 1996 | 95.19 | 1 | 58 | 3 | QKQH00000000 |
| PTA4_C2 | D | Caecum | 2.06 | 116 | 33,725 | 38.71 | 2021 | 95.12 | 1 | 64 | 3 | QKQG00000000 |
| PTA4_C4 | D | Caecum | 2.07 | 68 | 69,730 | 38.55 | 2050 | 95.13 | 1 | 61 | 3 | QKQF00000000 |
| PTA5_F1 | E | Faeces | 2.04 | 125 | 39,198 | 38.78 | 1954 | 95.20 | 1 | 64 | 3 | QKQE00000000 |
| PTA5_F4 | E | Faeces | 2.11 | 146 | 32,910 | 38.69 | 2052 | 95.27 | 1 | 61 | 3 | QKQD00000000 |
| PTA5_F11 | E | Faeces | 1.96 | 126 | 32,904 | 38.76 | 1909 | 95.15 | 1 | 63 | 3 | QKQC00000000 |
| PTA5_F13 | E | Faeces | 2.13 | 108 | 50,846 | 38.59 | 2144 | 95.07 | 1 | 60 | 3 | QKQB00000000 |
| PTA5_C4 | E | Caecum | 2.06 | 140 | 32,458 | 38.69 | 1995 | 95.04 | 1 | 64 | 3 | QKQA00000000 |
| PTA5_C5 | E | Caecum | 2.07 | 105 | 59,278 | 38.76 | 2007 | 95.27 | 1 | 58 | 3 | QKPZ00000000 |
| PTA5_C6B | E | Caecum | 2.71 | 657 | 22,978 | 38.7 | 2298 | 95.26 | 1 | 68 | 4 | QKPY00000000 |
| PTA5_C13 | E | Caecum | 2.19 | 158 | 34,750 | 38.84 | 2002 | 95.16 | 1 | 64 | 3 | QKPX00000000 |
| PTA6_C2 | F | Caecum | 2.23 | 191 | 34,786 | 38.86 | 2009 | 95.27 | 1 | 63 | 3 | QKPW00000000 |
| PTA6_C7 | F | Caecum | 2.19 | 154 | 39,778 | 38.74 | 2038 | 95.25 | 1 | 52 | 4 | QKPV00000000 |
| PTA6_F1 | F | Faeces | 2.28 | 219 | 31,238 | 38.88 | 2003 | 95.20 | 1 | 64 | 3 | QKPU00000000 |
| PTA6_F2 | F | Faeces | 2.17 | 147 | 45,484 | 38.59 | 2032 | 95.15 | 1 | 65 | 3 | QKPT00000000 |
| PTA6_F4 | F | Faeces | 2.57 | 442 | 40,200 | 38.57 | 2194 | 95.18 | 1 | 67 | 5 | QKPS00000000 |
| PTA6_F6 | F | Faeces | 2.47 | 414 | 26,145 | 38.74 | 2126 | 95.23 | 1 | 66 | 6 | QKPR00000000 |
| PTA6_F8 | F | Faeces | 2.04 | 135 | 39,822 | 38.76 | 1983 | 95.30 | 1 | 60 | 3 | QKPQ00000000 |

<sup>1</sup> > 500 bp
<sup>2</sup> Average Nucleotide Identity based on BLAST+ to the reference strain *Lb. reuteri* DSM20016

## Results and discussion

### *L. reuteri* isolation and typing

In the poultry GIT, *L. reuteri* forms biofilm in the crop and is among the most abundant *Lactobacillus* species commonly found in the caecum and colon. However, to date, only seven poultry *L. reuteri* strains (P43, An71, An166, 1366, JCM 1081, CSF8 and SKK-OGDONS-01) have been sequenced, and some of them included in previous phylogenetic analysis [2, 5, 36–38].

Here, by applying a colorimetric method on plate, a total of 70 reuterin-positive *L. reuteri* strains were isolated from poultry faeces and caecum from 6 poultry and from the crop obtained in the abattoir, and four reuterin-negative strains were selected. The identity of all strains was confirmed by sequencing of the 16s rRNA gene with 100 % similar to the reference strain *L. reuteri* DSM20016. Based on the similarity obtained from ERIC PCR profiles, 31 isolates with unique ERIC profiles were visually selected and further clustered with Bionumerics to confirm the uniqueness (Supplementary Figure S1). Based on the dendrogram obtained, 23 reuterin-positive with unique ERIC profiles and 2 reuterin-negative strains were identified for whole genome sequencing and further functional characterization. Among them, 13 were isolated from caecum and 10 from faeces while the 2 reuterin-negative strains originated from the crop. Considering the small sampling size, a high diversity of *L. reuteri* strains was found in this study. The colorimetric method applied allowed an easy phenotypic isolation of reuterin-positive colonies, that accounted for approximately 50 % of the bacterial colonies obtained on MRS isolation plates. This result indicated the high frequency of reuterin-producing *L. reuteri* strains in the poultry GIT, compared to the absence of reuterin production for rodent isolated strains [4].

Isolates from caecum did not appear to have a unique genetic profile when compared to isolates from faeces, and this could be due to the transition of strains from caecum to faeces, when not colonizers.

For the first time here, several *L. reuteri* strains were isolated from poultry and this will significantly enrich the database which will now account for 32 poultry strains making it 19 % of the total number of deposited *L. reuteri* strains, compared to 5 % before.

### Genomes analysis of *L. reuteri* poultry strains of this study

The whole genomes of 25 *L. reuteri* strains were sequenced and draft genomes were characterized. Strains had an average genome size of 2.159 ± 0.17 Mbp (Table 1). BUSCO assembly assessment showed good quality of assembly with 431 to 433 single-copy orthologous on a total of 433 for all assembled genomes (Supplementary Figure S2). Shotgun reads were assembled into contigs higher than 500 bp, ranging from 68 (PTA4_C4) to 657 (PTA5_C6B). The average N50 value, defined as the minimum contig length needed to cover 50 % of the genome, was 38337 ± 10.6 while the average guanine-cytosine (GC) content was 38.1 ± 0.11 %. The total number of coding sequences (CDs) ranged from 1909 to 2298, depending on the isolates. In every draft genome, 1 tmRNA gene, 45 to 68 tRNA genes and 3 to 6 rRNA genes were identified (Table 1). A total number of 2078 genes was annotated, with a core set of 817 shared genes. The number of annotated genes unique to each strain ranged from 0 to 31 (data not shown), showing a similar genetic content of the poultry isolates. The genomic features of *L. reuteri* isolates of this study in terms of genomes size, GC content and number of CDS were comparable with that of the 40 *L. reuteri* NCBI analyzed genomes from different hosts, suggesting not genetic diversity driven by the host (Supplementary Figure S3).

### Comparative phylogenetic analysis of *L. reuteri* strains from different hosts

The apparent relatedness between microbial community composition in the gut and host phylogeny has been interpreted as evidence of coevolution [39]. Symbiotic gut microbes associated with the host are predicted to evolve host-specific traits and, as a result, display enhanced ecological performances in their host [36, 40]. To assess evolution and adaptation of *L. reuteri* strains to different hosts, the gene content of poultry isolates was analyzed together with that of 40 *L. reuteri* strains available by NCBI, obtained from different hosts: human (6), rat (1), mouse (3), pig (4), sourdough (4), goat (5), sheep (4), cow (4), horse (3) and poultry (6). The gene content tree, in which strains sharing more genes clustered together, identified three main clusters namely cluster I, cluster II and cluster III, that contained previously observed *L. reuteri* lineages [2]. Those host-adapted lineages were first described after the characterization of the genetic structure of *L. reuteri* strains isolated from human, mouse, rat, pig, chicken and turkey, and the same lineage names were also applied in our study for coherency [2]. Cluster I, corresponding to the previously defined poultry/human lineage IV, comprised all 25 *L. reuteri* poultry isolates of this study and all, except one (P43), poultry NCBI isolates. The same cluster also included two humans strains (SD2112 and CF48-3A) and this was also the case in all previous phylogenetic analysis of *L. reuteri* isolates from different hosts [2, 36, 38]. The fact that even with a much higher number of poultry isolates the two human isolates, respectively isolated from human breast milk in Peru (SD2112) and from the child faeces in Finland (CF48-3A), still cluster together with 29 poultry isolates, suggests that these strains could be of poultry origin. In a previous study, administering human isolates of lineage VI were shown to colonize the poultry GIT and therefore may be a necessary colonization factors indicating rather co-evolution events of human and poultry isolates [36]. In our study, cluster II included the majority of herbivorous isolates (new defined herbivorous lineage VII of our study) in addition to four human (DSM20016, MM2_3, IRT and JCM1112), one sourdough (CRL1098), one rodent (mlc3) and one porcine (20-02) strains, belonging to lineages II, III and IV previously defined [2]. Cluster III was composed of pig, herbivorous, sourdough and rodent isolates (Figure 1) corresponding to lineages I, III, V and newly defined in this study: herbivorous VIII. ANI analysis of the 65 *L. reuteri* isolates (Supplementary Figure S4) identified the same three clusters, with exception of published strains 20_02 and mlc3 that were assigned to cluster III instead of cluster II.

**Figure 1.**
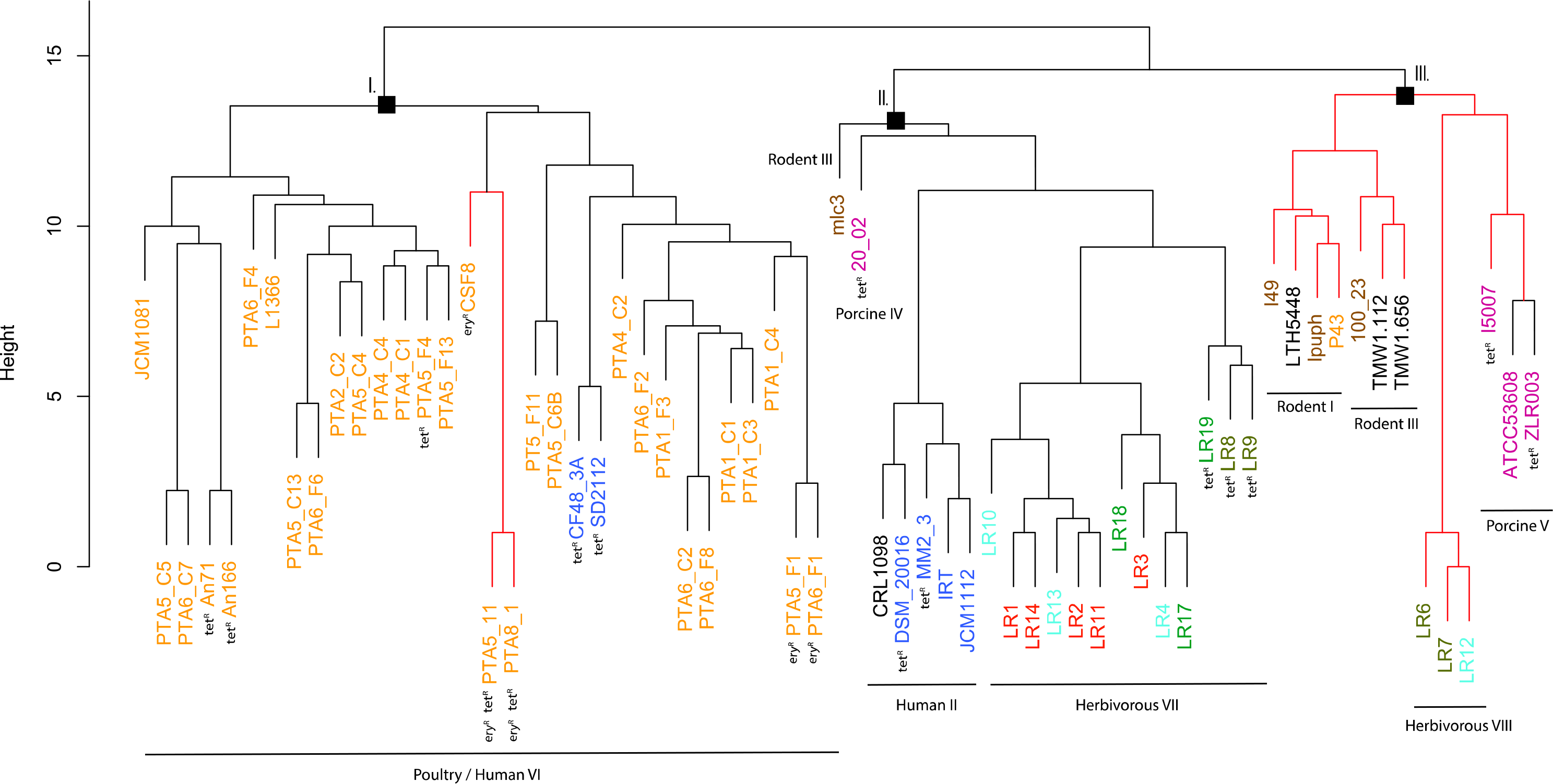
Phylogenetic tree based on gene content matrix (presence or absence of annotated gene) of 65 *L. reuteri* strains from different hosts (25 genomes from this study and 40 from NCBI, Table S1). Different colours represent different hosts, blue: human; yellow: poultry; pink: pig; brown: mouse/rat; light blue: cow; red: goat; green: horse; brown/green: sheep; black: sourdough. The red branches indicate the reuterin-negative strains. I, II and III indicate identified clusters. Strains which harbour AMR genes for tetracycline (tet) and erythromycin (ery) are indicated as tetR and eryR, respectively.

The phylogenetic tree based on core genomes of the 65 genomes covered a core of 1152 genes per genome, for a total of 74880 genes. In agreement with the gene content tree, all isolated *L. reuteri* poultry strains clustered together with NCBI poultry isolates (strains JCM1081, CSF8, An71 and An166), except P43, forming poultry/human lineage VI [2]. As indicated above, this cluster also included the two human isolates CF48-3A and SD2112 (cluster I, Figure 2). Cluster II was composed of strains from human lineage II and herbivorous lineage VII, as previously presented in gene content tree, and the same applied for cluster III which was composed of porcine lineage V and herbivorous lineage VIII. *L. reuteri* strains belonging to rodent lineage III, rodent lineage I and porcine lineage IV clustered differently from the gene content tree, which includes accessory genes, indicating gene loss or acquisition of genes by horizontal genetic transfer.

**Figure 2.**
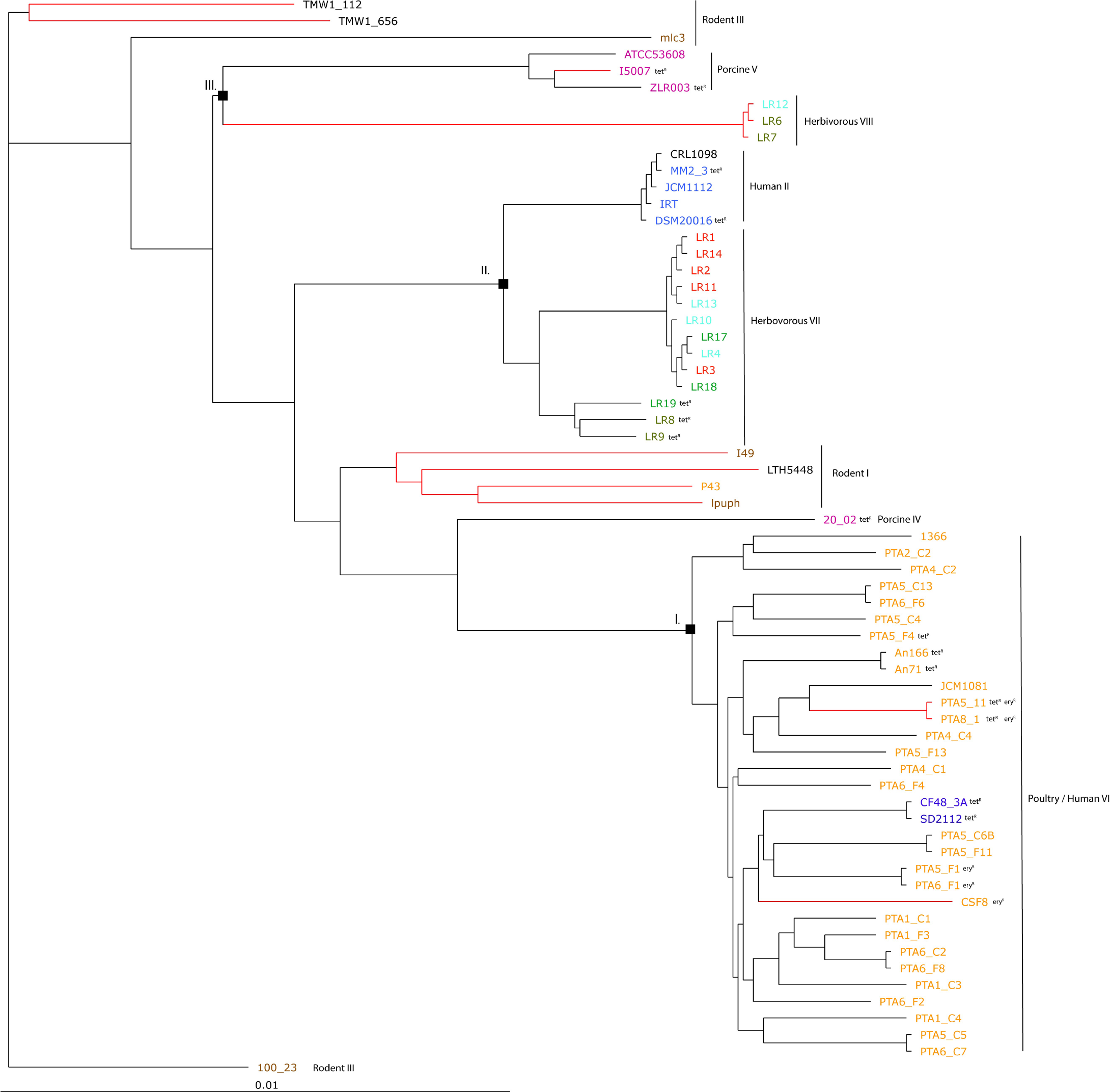
Phylogenetic tree based on core genes of 65 *L. reuteri* strains from different hosts. **(25 genomes from this study and 40 from NCBI, Table S1)** based on neighbour joining method (PHYLIP implementation). Different colours represent different hosts, blue: human; yellow: poultry; pink: pig; brown: mouse/rat; light blue: cow; red: goat; green: horse; brown/green: sheep; black: sourdough. The red branches indicate the reuterin-negative strains. I, II and III indicate identified clusters in the gene content tree (Figure 1). Strains which harbour AMR genes for tetracycline (tet) and erythromycin (ery) are indicated as tetR and eryR, respectively.

Overall, all three-different analysis (gene content tree, ANI analysis and core genome tree) confirmed the existence and composition of the poultry/human lineage VI, which was substantially enriched by the 25 poultry isolates from our study. Those strains appear to share both core and accessory genes and to be highly similar also at nucleotide level.

Here, for the first time, lineages of herbivorous strains were defined. Besides individual lineages with isolates from different hosts, the cluster analysis performed here grouped together human and poultry lineage VI strains (cluster I), human lineage II with herbivorous lineage VII strains (cluster II) and porcine lineage V with herbivorous lineage VIII strains (cluster III), suggesting possible co evolutions of those three clusters.

### Genes unique for the poultry/human lineage VI

Presence of host specific lineages in itself does not necessary provide evidence for natural selection, as a cluster can arise by neutral processes, such as genetic diversion [2]. It has been demonstrated how strains from rodent display elevated fitness in mice, and biofilm formation in the forestomach is restricted to strains from rodent lineages. Moreover, *L. reuteri* rodents strains were able to effectively colonize rodent host in vivo [2, 5, 36]. However, this was not the case for pig isolates [36, 37]. Here, forty unique genes of the poultry/human lineage VI were identified (Supplementary Table S3) and were mainly categorized as transport proteins DNA-binding proteins and transferase proteins. Such genes could not be directly linked with adaptation to poultry physiology or feeding (Figure 3). Further studies are needed to elucidate the specific role (s) of those unique genes that may be linked to poultry adaptation.

**Figure 3.**
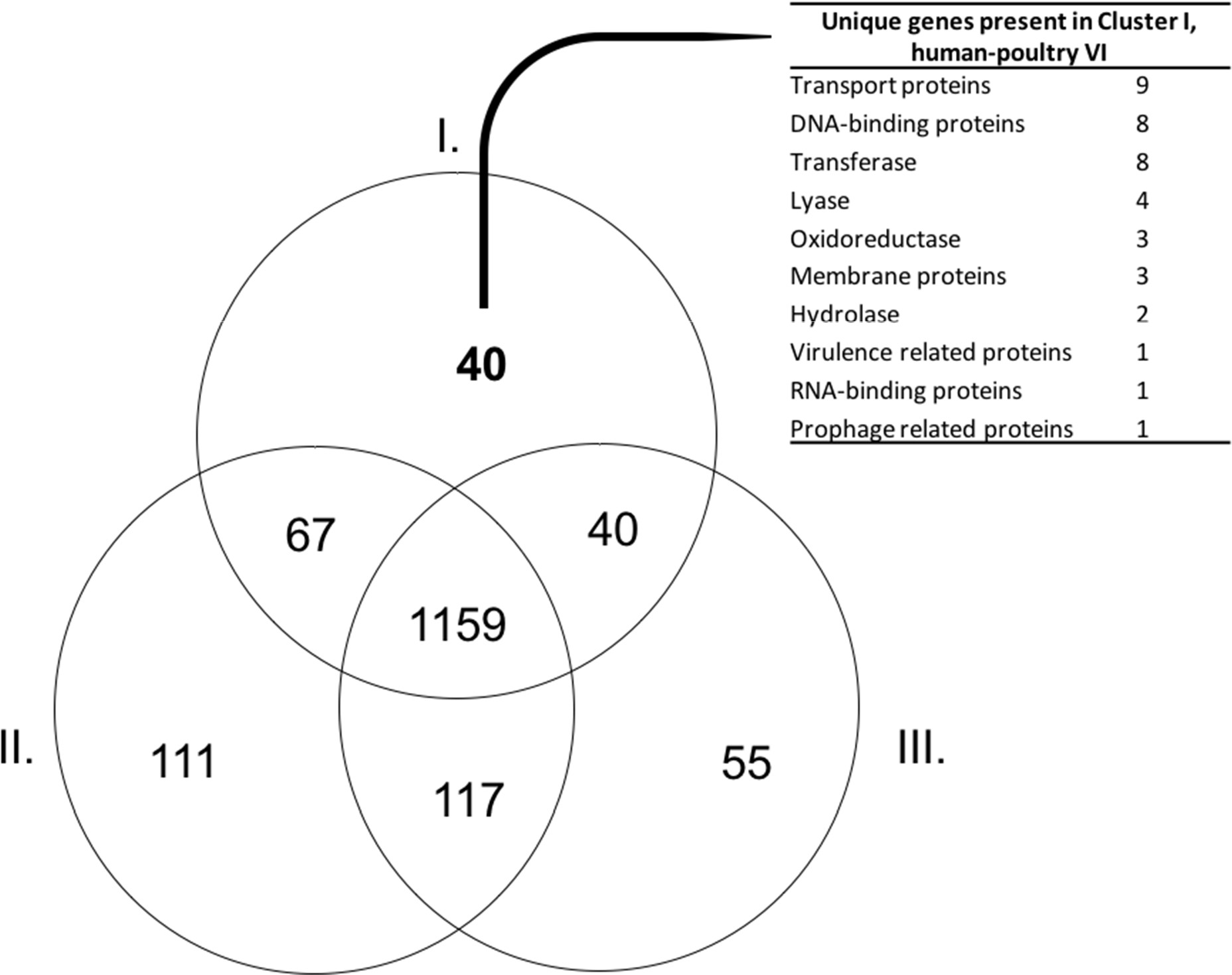
Venn diagram showing the number of shared genes of the core genomes of cluster I (poultry/human lineage VI), cluster II (lineages human II, vertebrate II, porcine IV and rodent III) and cluster III (lineages rodent I, rodent III, vertebrate I and porcine V). Major functional classes unique in cluster I are shown in the table. Individual genes are listed in Supplementary Table S3.

### Reuterin synthesis

The presence and composition of reuterin operon genes (*pdu-cbi-cob-hem*) was investigated in all 65 genomes (Figure 4) and reuterin production was determined as a marker for PduCDE activity. All *L. reuteri* strains isolated in this study that possessed the complete *pdu-cbi-cob-hem* operon produced 3-HPA when incubated in 600 mM glycerol. In contrast strains PTA5_11 and PTA8_1 only possessed *hemH, hemA, cobC* and *cobB* but lacked all the others operon genes (Figure 4) and therefore did not produce reuterin. Under the conditions of the test, 3-HPA yield ranged from 156.9 mM ± 11.0 (PT6_F1) to 330.2 mM ± 14.9 (PTA4_C4) starting from 600 mM glycerol (Figure 2). Reference strains DSM 20016 (human lineage II) produced 132.8 ± 4.3 mM while SD2112 (human lineage VI) produced 432.9 ± 9.0 mM.

**Figure 4.**
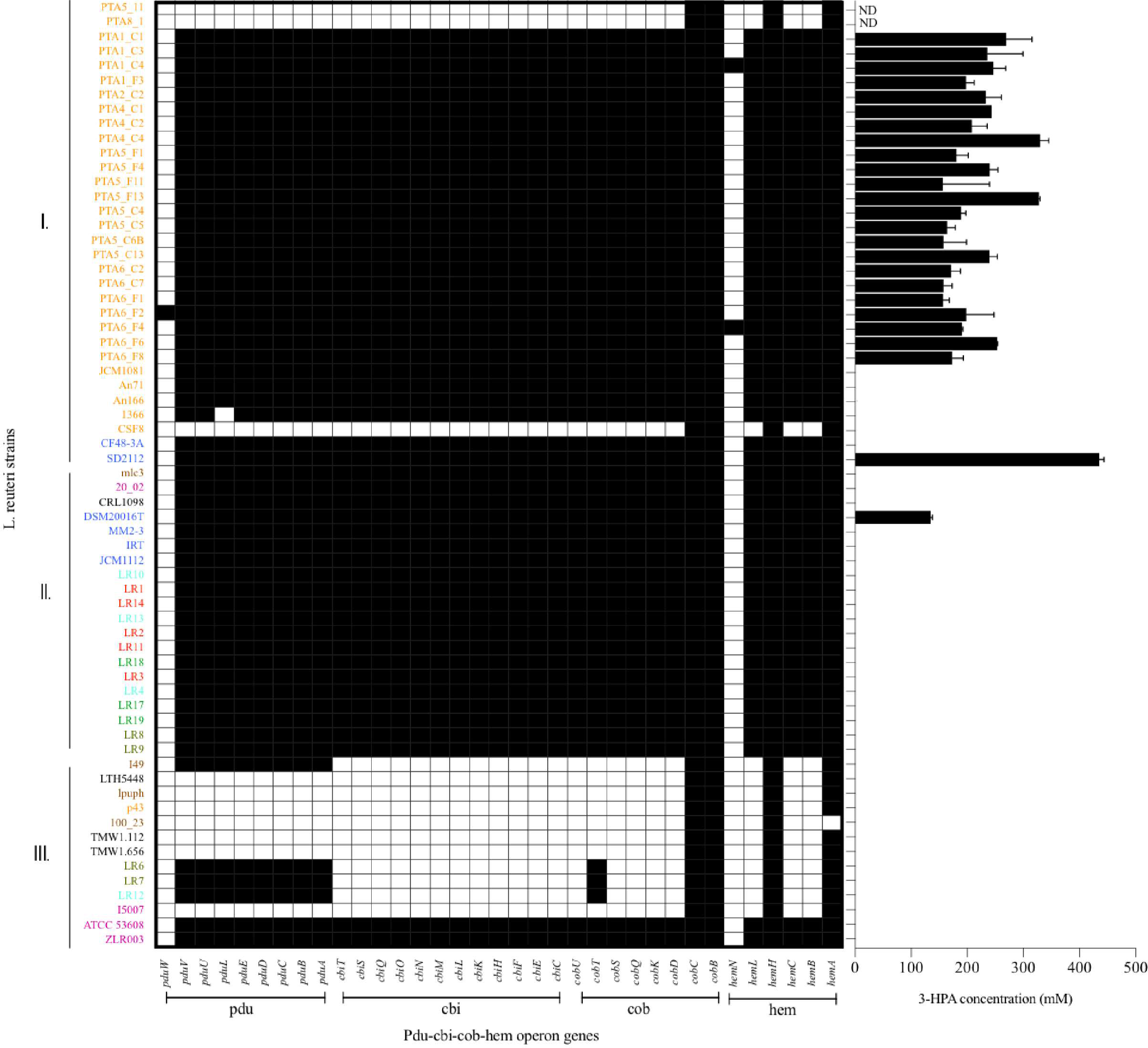
Reuterin operon genes (*pdu-cbi-cob-hem*) detected in the genomes of 65 *L. reuteri* listed in Table S1 and 3-HPA production of 25 strains isolated from poultry in this study. Black indicates presence of a gene. Different colours represent different hosts, blue: human; yellow: poultry; pink: pig; brown: mouse/rat; light blue: cow; red: goat; green: horse; brown/green: sheep; black: sourdough. ND not determined.

Cluster I strains (corresponding to poultry/human lineage VI) all harbored *pdu-cob-cbi-hem* genes, except for a small sub-cluster of three by reuterin-negative poultry isolates CSF8, PTA5_11 and PTA8_1. Isolates assigned to Cluster II possessed *pdu-cob-cbi-hem* and have been shown to mostly form reuterin on MRS agar plates overlaid with 500mM glycerol agar [4]. Isolates of this cluster lacked *pduW* and *hemN* genes, which therefore seem not essential for reuterin production. These genes were also not detected in the vast majority of reuterin-positive strains isolated in our study (Figure 4). In contrast, the prevalence of *pdu-cob-cbi-hem* scattered in Cluster III comprising isolates of rodent lineages I and III, herbivorous and porcine lineages VIII and V, respectively. Only strains ATCC53608 and ZLR003 possessed a complete functional *pdu-cob-cbi-hem* operon while cluster III rodent isolates lpuph and 100_23 lack the majority of the operon genes.

Interestingly, herbivore isolates LR6, LR7, LR12 and the rodent isolate I49 possessed the *pdu* but not the *cbi* and *cob* genes while cobalamin is a cofactor for 3-HPA production. Therefore these strains are likely not able to form 3-HPA from glycerol (or propanal from 1,2-propanediol) and are reuterin-negative (indicated with red branches in Figure 1 and Figure 2) unless they acquire the vitamin from other sources or microbes. Rodent strains, which are mostly reuterin-negative based on the analysis of operon genes in this study, have been previously suggested as being at the root of the evolutionary history of *L. reuteri*-host associates [36]. This data suggests that the *pdu-cbi-cob-hem* operon and thus reuterin production was acquired later during evolution by *L. reuteri* strains in rodents and also in poultry/human lineage VI strains.

### Antimicrobial sensitivity profiles of *L. reuteri* strains

The horizontal transfer of AMR genes is a rising risk concern, and the absence of transferable AMR genes must be demonstrated for application of new strains in food and feed (WHO, 2017). Antimicrobials used in farmed animals for diseases prevention have been associated with an increase of frequency of resistant bacteria in chickens, swine, and other food-producing animals GIT [41]. The high use of antimicrobials in animal production is likely to accelerate the development of antimicrobial resistance in pathogens, as well as in commensal organisms, resulting in treatment failures, economic losses and source of gene pool for transmission to humans [41]. Poultry is one of the most widespread food industries worldwide and various antimicrobials are used to treat infections mainly in young poultry [42, 43].

The antimicrobial susceptibility profile of *L. reuteri* poultry isolates and reference strains DSM20016 and SD2112 showed that all strains were sensitive to cefotaxime (MIC values from 0.016 to 1 μg/mL). All poultry isolates were also sensitive to penicillin with MIC values from 0.02 and 3 μg/mL, in contrast to DSM20116 and SD2112 that showed resistance to this antibiotic (MIC> 256 ug/mL). Penicillin resistance were shown to result from point mutations of the chromosomally located genes encoding penicillin-binding proteins Pbp [44]. Penicillin-binding genes *pbpX, pbpF, pbpB* and *ponA* were identified in all 65 strains with both resistant and sensitive phenotype. Several SNPs were observed for DSM20016 (Table 3), especially in *ponA* and *pbpX_2* which led to point mutations of the corresponding proteins. Only one amino acid substitution at position 134 of PbpX_2 was shared among the two resistant strains: DSM20016 possessed a Q instead of H (H_134_Q) while in the same position strain SD2112 had a Y (H_134_Y). The substitution at this position may be the one responsible for the penicillin resistant phenotype observed for DSM20016 and SD2112.

**Table 2.**
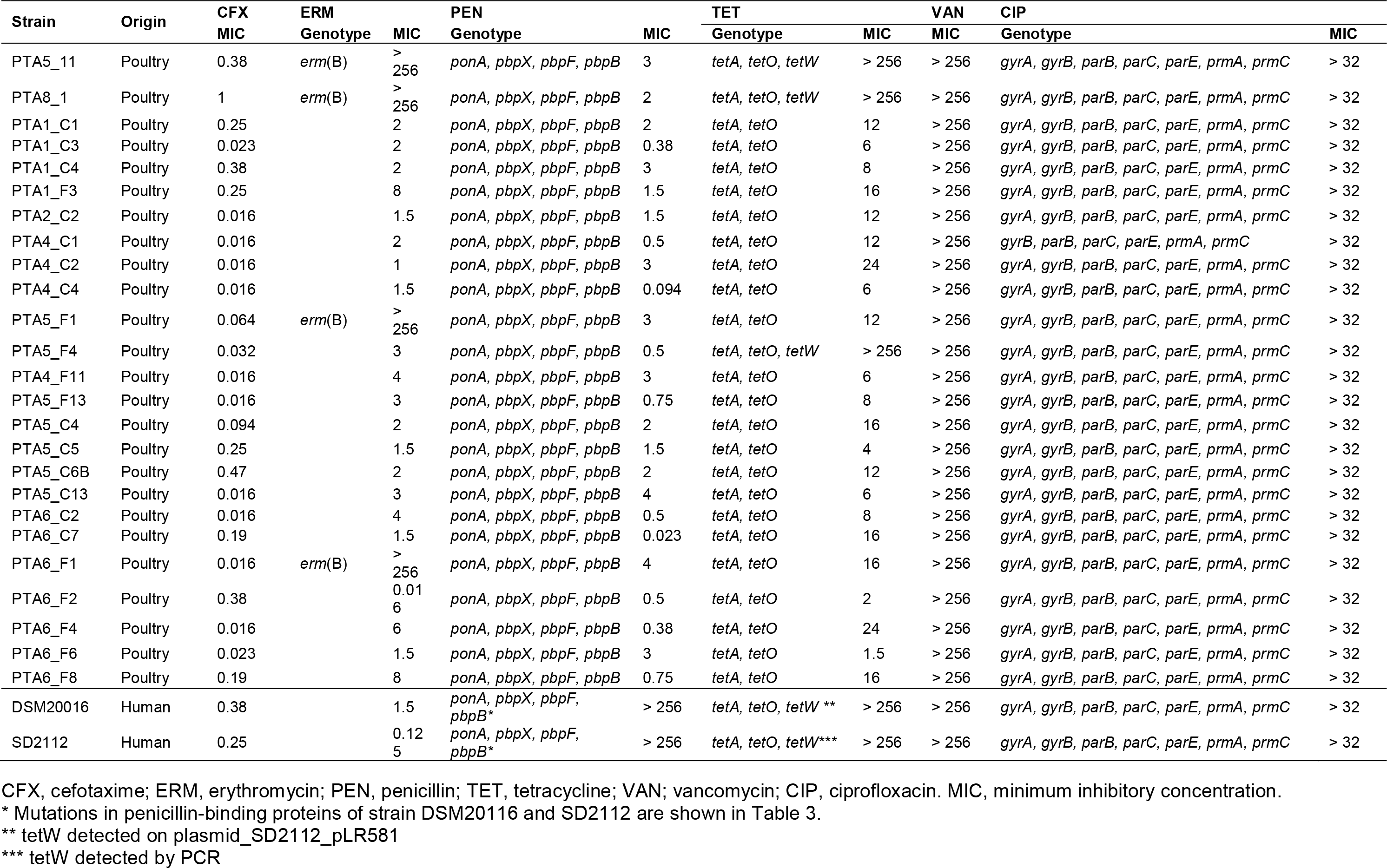
Antimicrobial resistance (AMR) profiles of 25 *L. reuteri* strains isolated from poultry in this study and of reference strains measured using MTS™ strips (MIC, ug/mL), and associated AMR genes detected in their draft genomes.

**Table 3.** Point mutations identified in penicillin-binding protein genes of resistant strains *L. reuteri* DSM20016 and *L. reuteri* SD2112. Amino acids changes in the resistant strains, compared to all other 25 *L. reuteri* sensitive strains, are indicated.

| Strain | Gene | Mutation(s) in Penicillin-binding proteins |
| --- | --- | --- |
| DSM20016 | *ponA* | D<sub>322</sub>N / S<sub>537</sub>A / V<sub>538</sub>A / D<sub>712</sub>E/ A<sub>723</sub>V/ S<sub>725</sub>N/ A<sub>740</sub>T/ D<sub>411</sub>V/ Y<sub>412</sub>F / Q<sub>491</sub>L |
|  | *pbpB* | A<sub>361</sub>T / H<sub>495</sub>Q / T<sub>586</sub>P / V<sub>613</sub>A / L<sub>714</sub>S |
|  | *pbpF* | D<sub>9</sub>G / T<sub>22</sub>A |
|  | *pbpX_1* | V<sub>26</sub>I / T<sub>61</sub>P / I<sub>62</sub>V / A<sub>63</sub>T / K<sub>100</sub>R / D<sub>123</sub>E |
|  | *pbpX_2* | V<sub>19</sub>I / L<sub>26</sub>V / H<sub>30</sub>R / I<sub>49</sub>L / V<sub>86</sub>I / D<sub>96</sub>N / T<sub>111</sub>A / N<sub>127</sub>T / K<sub>132</sub>R / **H<sub>134</sub>Q** / R<sub>141</sub>H / T<sub>170</sub>A / R<sub>185</sub>N / Q<sub>187</sub>H / K<sub>244</sub>N / T<sub>252</sub>I / L<sub>255</sub>F / Y<sub>320</sub>F / D<sub>322</sub>A / K<sub>333</sub>T |
| SD2112 | *ponA* | D<sub>411</sub>V / Y<sub>412</sub>F / Q<sub>491</sub>L |
|  | *pbpX_1* | R<sub>141</sub>H |
|  | *pbpX_2* | E<sub>124</sub>K / **H<sub>134</sub>Y** |
|  | *pbpX_3* | P<sub>57</sub>L / L<sub>193</sub>P |

In bold, a shared mutation for both resistance strain with a different amino acids substitution.

Four poultry isolates of this study (PTA5_11, PTA8_1, PTA5_F1 and PTA6_F1) showed resistance phenotype (MIC > 256 μg/mL) to erythromycin, confirmed by the presence of *ermB*, which is usually found on a plasmid [45, 46]. The *ermB* gene encodes enzymes that modify the 23S rRNA by adding one or two methyl groups, reducing the binding to the ribosome of different classes of antibiotics [47]. The presence of *ermB* in the genome of resistant strains was confirmed by using PCR (data not shown). Among the 40 NCBI strains analyzed in this study, *ermB* was also detected in the genome of poultry isolate CSF8 (Figure 1 and Figure 2). Erythromycin resistance has been found in *Bacillus cereus, Pseudomonas aeruginosa, C. jejuni, C. coli, Clostridium perfringens* and *Enterococcus* poultry isolates [43].

Tetracycline resistance genes tetA and tetO were detected in all 65 *L. reuteri* genomes analyzed, but did not appear to be directly correlated with this resistance phenotype (Table 2 and Supplementary Table S2). The only 3 poultry isolates resistant to tetracycline (MIC > 256 μg/mL) possessed additionally the *tetW* in their genome (PTA5_11, PTA8_1 and PTA5_F4), similar to DSM20016 and SD2112 (MIC > 256 μg/mL). Tetracycline resistance has been described to be allocated on a plasmid for example for SD2112 (pLR581) [34]. For DSM20016, *tetW* was not detected on the chromosome but was amplified by PCR. This suggest that *tetW* is likewise located on a plasmid (data not shown). Besides *tetW, tetM, tetL* and *tetC* also associated with tetracycline resistance were identified in the genomes of different isolates (MM2_3, I5007, 20_02, ZLR003, LR8, LR9 and LR19). Interestingly, tetracycline-resistant poultry isolates (An71, An166) harboured *tetW* while herbivorous resistant strains (LR8, LR9 and LR19) possessed *tetM*. In contrast, pig isolates harboured *tetM* and *tetW* (I5007 and 20_02) or *tetW* and *tetL* (ZLR003) (Supplementary Table S2).

Tetracyclines are widespread antimicrobials extensively used in livestock [48, 49]. Tetracyclines are a family of compounds frequently employed due to their broad spectrum of activity as well as their low cost, compared with other antimicrobials [48]. However, *in vivo* transferability of *tetW* from *L. reuteri* ATCC55730 to other human gut microbes in a double-blind clinical study was not demonstrated [50].

Lactobacilli are suggested to be intrinsically resistant to vancomycin and ciprofloxacin [51]. In agreement with previous studies [15], all poultry *L. reuteri* strains of this study were resistant to vancomycin (MIC > 256 μg/mL). Vancomycin resistance in lactobacilli has been shown to be linked to the *vanX* gene encoding a d-Ala-d-Ala dipeptidase [52]. Other vancomycin resistance genes were described in the literature with *vanA*, *vanB*, *vanC* and *vanE* [34, 52]. None of those genes were detected in the genomes of *L. reuteri* in agreement with previous studies [34, 52]. However, changes in membrane composition have also been associated with intrinsic vancomycin resistance [53]. *vanH*, a D-lactate dehydrogenase gene was detected in one pig strain (ATCC 53608) [54]. The same gene had been previously associated with vancomycin resistance of *Enterococcus faecium* [55]. Ciprofloxacin resistance seems to be widely spread among lactobacilli [51, 56, 57]. All 25 poultry isolates were resistant to ciprofloxacin with MIC >32 μg/mL, and all 65 genomes analysed possessed the six genes for which mutations were correlated with ciprofloxacin resistance, namely *gryB, parB, parC, parE, prmA* and *prmC*, while gryA was present in all but one (PTA4_C1) isolate.

*L. reuteri* has been affiliated to different hosts, which might be exposed to different levels and types of antimicrobials and is commercially used as probiotic in food and feed [11, 12, 58, 59]. The results of this study indicated that *L. reuteri* poultry isolates harbour some AMR genes. In view of application *L. reuteri* in feed to prevent pathogen infection, strains without transferable AMR genes must be carefully selected. The first applied *L. reuteri* probiotic strains (SD2112) harbours the tetracycline resistant gene *tetW* on a plasmid [34]. This strain was however curated for the plasmid free daughter strain DSM17938 [44], which is used for commercially.

## Conclusion

In conclusion, this study substantially enriched the pool of poultry *L. reuteri* strains for comparative genomic and evolutionary studies. The phylogenetic analysis confirmed and straightened the co-evolution of human isolates of lineage VI with poultry (poultry/human lineage VI). However, due to the high number of poultry isolates in this lineage compared to only two human isolates, we speculate a possible cross contamination during isolation of the two human strains belonging to lineage VI. The pool of *L. reuteri* poultry isolates of this study may be useful to select and characterize high potential strains exhibiting reuterin activity, and develop application in poultry, as a natural antimicrobial system to prevent pathogen infections and colonization of the poultry GIT.

## Acknowledgments

This project was funded by COOP Research Program on “Sustainability in Food Value Chains” of the ETH-Zürich World Food System Centre and the ETH Zürich Foundation for supporting this project. The COOP Research Program is supported by the COOP Sustainability Fund. Paul Tetteh Asare was supported by the Swiss Government Excellence PhD Scholarships for foreign students. The authors are grateful to Alfonso Die for the HPLC measurements

The EDGAR platform used in this study is financially supported by the BMBF grant FKZ 031A533 within the de.NBI network. Data analysed in this paper were generated in collaboration with the Genetic Diversity Centre (GDC), ETH Zurich.

