## Supplementary Figure S1 for "Isolation and comparative genomic analysis of reuterin-producing *Lactobacillus reuteri* from poultry gastrointestinal tract"

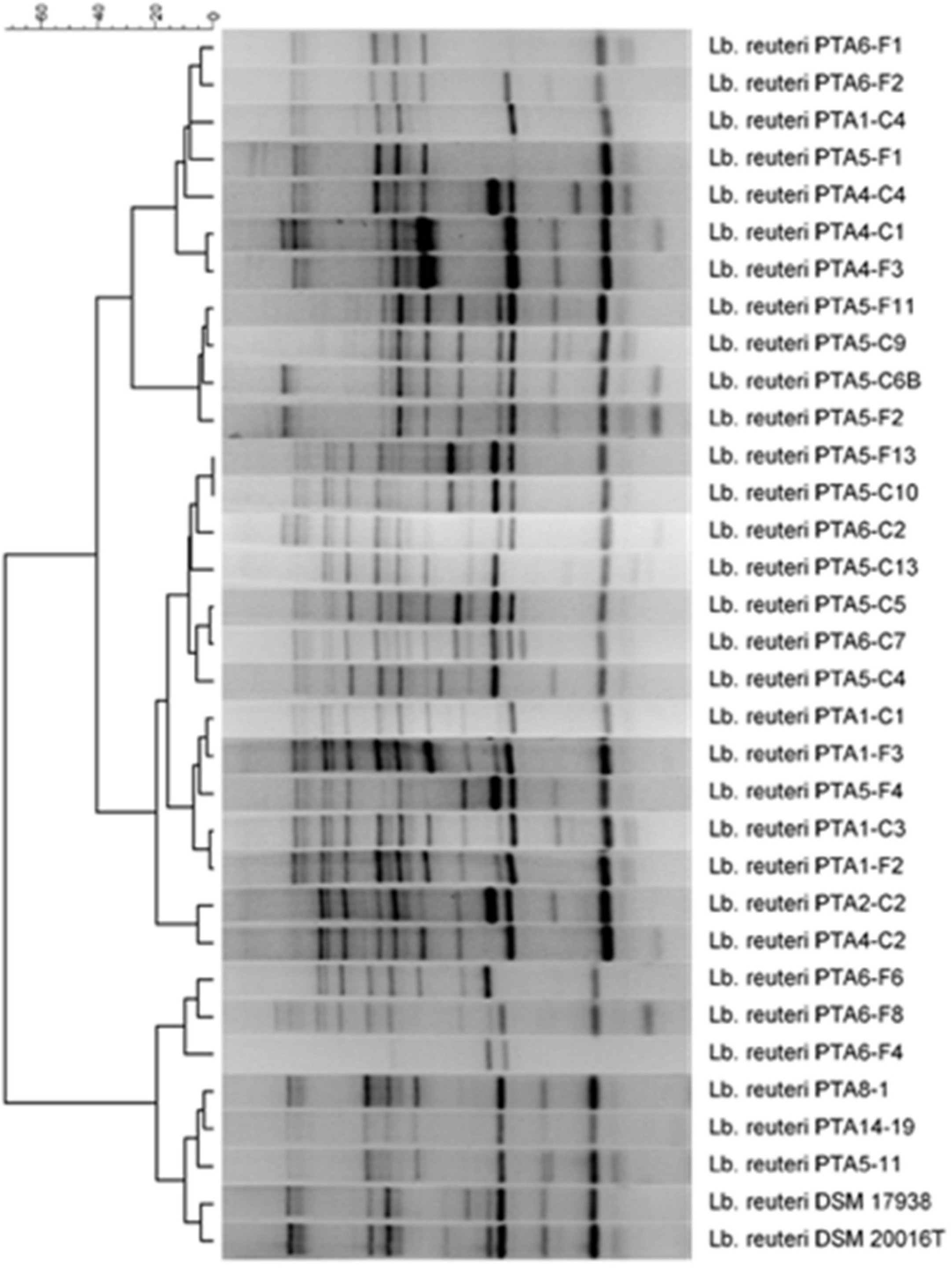


**Supplementary Figure S1**. ERIC PCR profiles of selected *L. reuteri* isolated in this study from poultry faeces and caecum.
