## Supplementary Figure S2 for "Isolation and comparative genomic analysis of reuterin-producing *Lactobacillus reuteri* from poultry gastrointestinal tract"

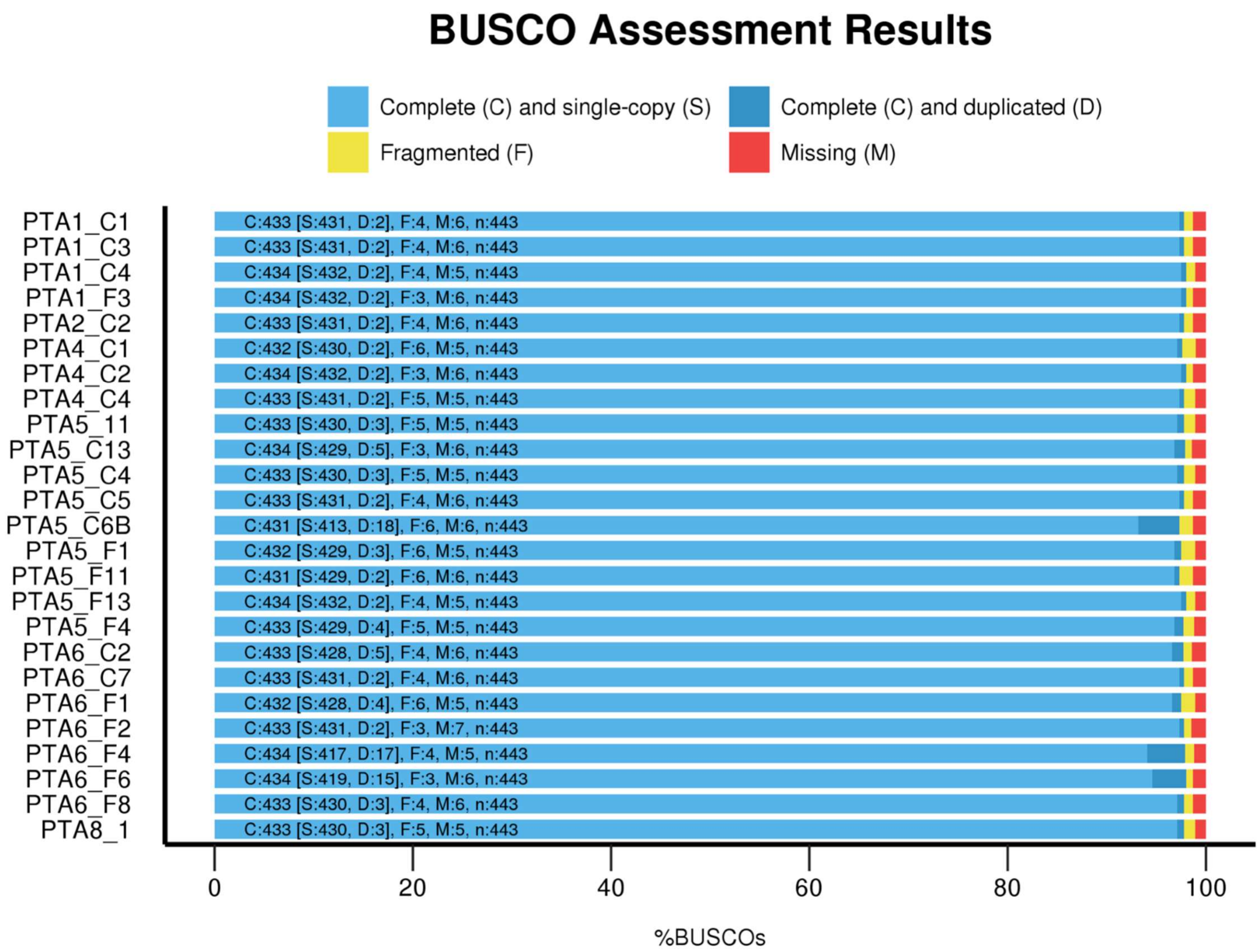


**Supplementary Figure S2.** BUSCO genome assembly assessment of 25 draft genomes of *L. reuteri* poultry isolates from this study
