## Supplementary Figure S3 for "Isolation and comparative genomic analysis of reuterin-producing *Lactobacillus reuteri* from poultry gastrointestinal tract"

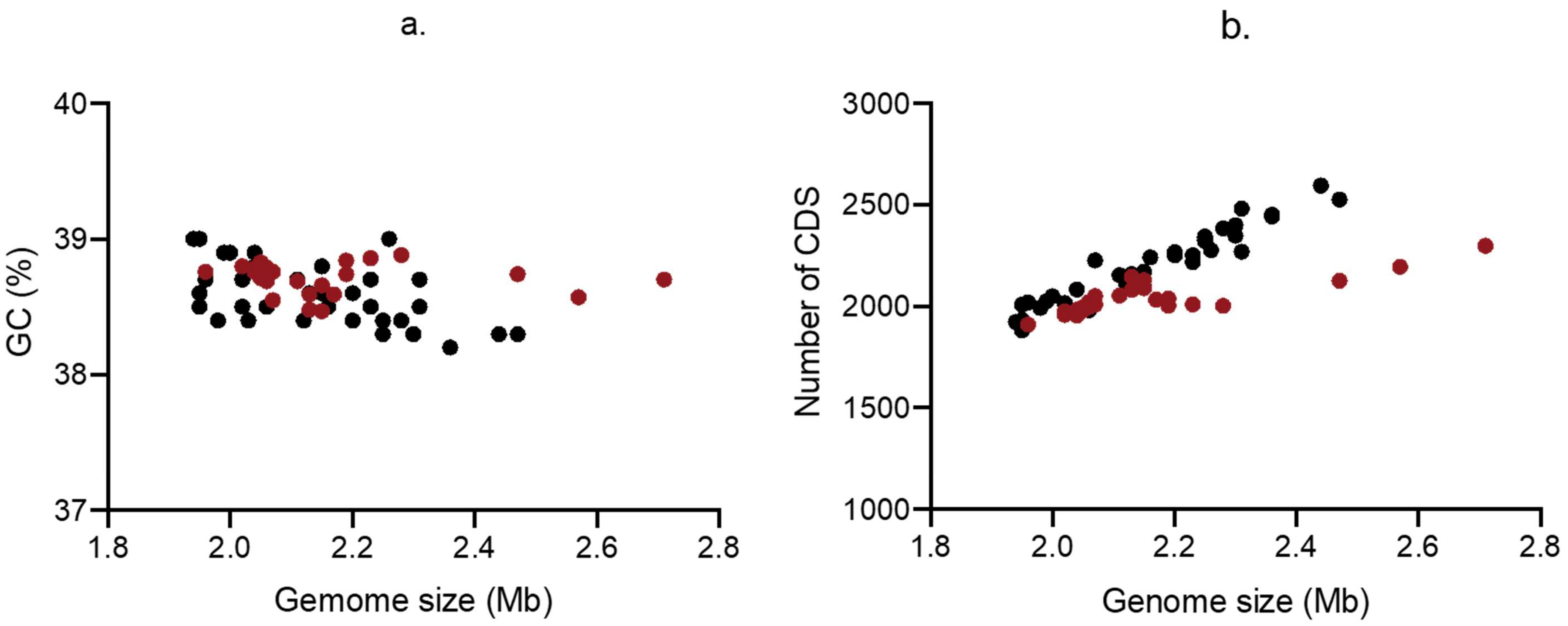


**Supplementary Figure S3.** Genomic characteristics of *L. reuteri* poultry isolates of this study (red dots) compared to 40 NCBI *L. reuteri* deposited genomes (black dots); a) correlation between GC content (%) and genome size; b) correlation between genome size and the number of coding sequences (CDSs).
