## Supplementary Figure S4 for "Isolation and comparative genomic analysis of reuterin-producing *Lactobacillus reuteri* from poultry gastrointestinal tract"

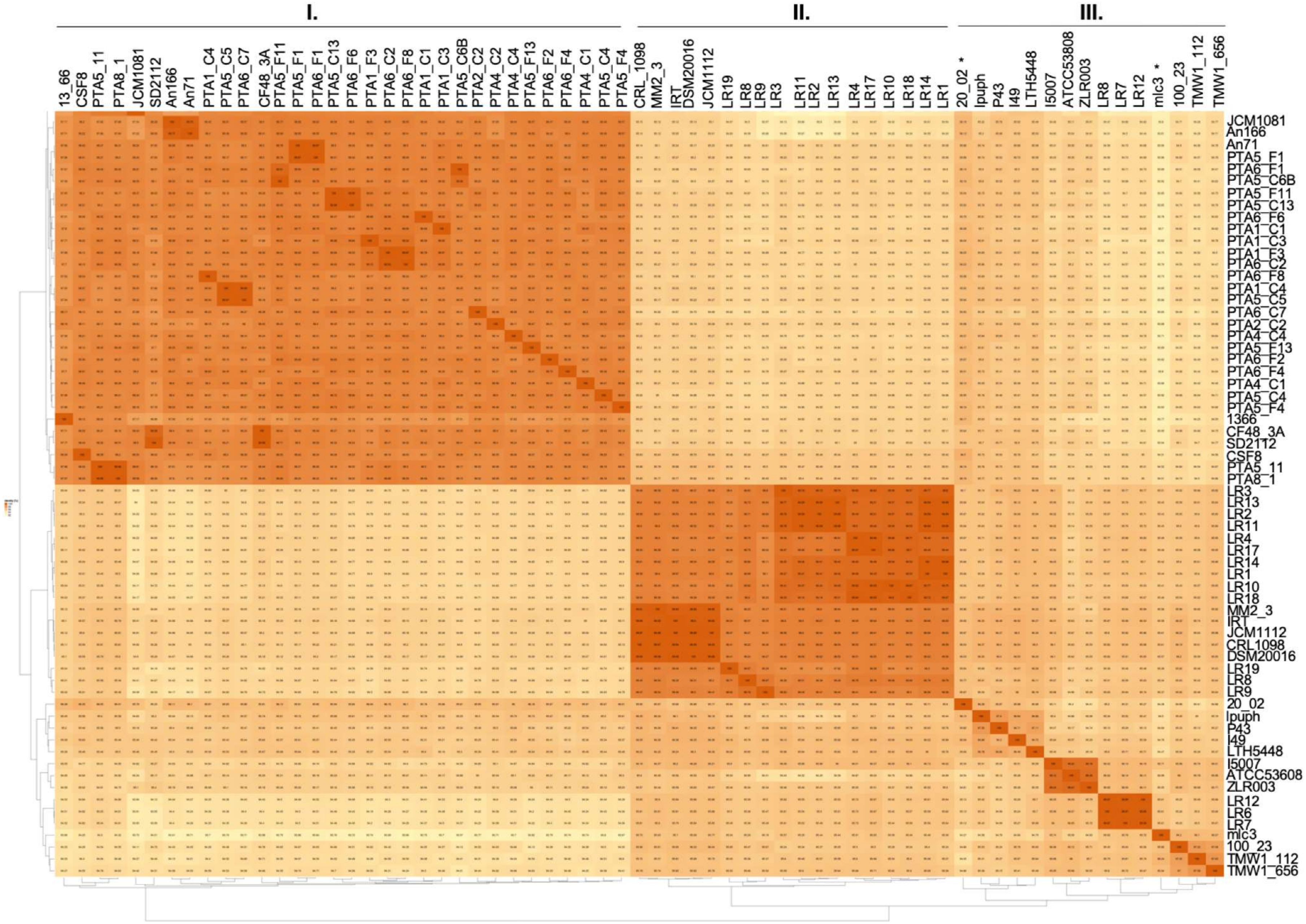


**Supplementary Figure S4.** Average nucleotide identity (ANI) of *L. reuteri* isolates calculated with EDGAR 2.3. Asterisk (*) indicate two strains belonging to cluster II in the genetic tree
