## Supplementary Table S1 for "Isolation and comparative genomic analysis of reuterin-producing *Lactobacillus reuteri* from poultry gastrointestinal tract"

**Supplementary Table S1.** *L. reuteri* strains with genomes published in NCBI used for comparative genomics. Genomes were retrieved from NCBI and annotated with the same pipeline used for the 25 draft genomes of this study.

| *L. reuteri* strains | Host | Country of Origin | NCBI accession number |
|---|---|---|---|
| P43 | Poultry | USA | MCNS00000000 |
| An71 | Poultry | N.D. | NZ_NFHN00000000 |
| An166 | Poultry | N.D. | NZ_NFKV00000000 |
| 1366 | Poultry | Denmark | NZ_NBBG00000000 |
| JCM 1081 | Poultry | Japan | NZ_NBBD00000000 |
| CSF8 | Poultry | USA | NZ_NBBE00000000 |
| DSM20016 | Human | Germany | NC_009513 |
| IRT | Human | South Korea | NZ_CP011024 |
| JCM 1112 | Human | Japan | NC_010609 |
| CF48_3A | Human | USA | NZ_ACHG00000000.1 |
| MM2_3 | Human | USA | NZ_ACLB00000000.1 |
| SD2112 | Human | Peru | NC_015697 |
| 100_23 | Rat | New Zealand | AAPZ00000000 |
| I49 | Mouse | Switzerland | NZ_CP015408 |
| mlc3 | Mouse | USA | AEAW00000000 |
| lpuph | Mouse | USA | AEAX00000000 |
| ATCC 53608 | Pig | Sweden | NZ_LN906634 |
| I5007 | Pig | China | NC_021494 |
| ZLR003 | Pig | China | NZ_CP014786 |
| 20_02 | Pig | Germany | CZDD00000000.1 |
| CRL1098 | Sourdough | Argentina | LYWI00000000 |
| TMW1.656 | Sourdough | Germany | JOSW00000000 |
| TMW1.112 | Sourdough | Germany | JOKX00000000 |
| LTH5448 | Sourdough | Germany | JOOG00000000 |
| LR1 | Goat | China | QGID00000000 |
| LR2 | Goat | China | QGIC00000000 |
| LR3 | Goat | China | QGIB00000000 |
| LR11 | Goat | China | QGIA00000000 |
| LR14 | Goat | China | QGHZ00000000 |
| LR6 | Sheep | China | QGHY00000000 |
| LR7 | Sheep | China | QGHX00000000 |
| LR8 | Sheep | China | QGHW00000000 |
| LR9 | Sheep | China | QGHV00000000 |
| LR4 | Cow | China | QGHU00000000 |
| LR10 | Cow | China | QGHT00000000 |
| LR12 | Cow | China | QGHS00000000 |
| LR13 | Cow | China | QGHR00000000 |
| LR17 | Horse | China | QGHQ00000000 |
| LR18 | Horse | China | QGHP00000000 |
| LR19 | Horse | China | QGHO00000000 |
