## Supplementary Table S2 for "Isolation and comparative genomic analysis of reuterin-producing *Lactobacillus reuteri* from poultry gastrointestinal tract"

**Supplementary Table S2.** AMR genes detected in the genomes of *L. reuteri* strains isolated from different hosts and published in NCBI.

| Strain | Origin | ERM | PEN | TET | VAN | CIP |
|---|---|---|---|---|---|---|
| P43 | Poultry | | *ponA, pbpX, pbpF, pbpB* | *tetA, tetO* | | *gyrA, gyrB, parB, parC, parE, prmA, prmC* |
| An71 | Poultry | | *ponA, pbpX, pbpF, pbpB* | *tetA, tetO, tetW* | | *gyrA, gyrB, parB, parC, parE, prmA, prmC* |
| An166 | Poultry | | *ponA, pbpX, pbpF, pbpB* | *tetA, tetO, tetW* | | *gyrA, gyrB, parB, parC, parE, prmA, prmC* |
| 1366 | Poultry | | *ponA, pbpX, pbpF, pbpB* | *tetA, tetO* | | *gyrA, gyrB, parB, parC, parE, prmA, prmC* |
| JCM1081 | Poultry | | *ponA, pbpX, pbpF, pbpB* | *tetA, tetO* | | *gyrA, gyrB, parB, parC, parE, prmA, prmC* |
| CSF8 | Poultry | *erm*(B) | *ponA, pbpX, pbpF, pbpB* | *tetA, tetO* | | *gyrA, gyrB, parB, parC, parE, prmA, prmC* |
| IRT | Human | | *ponA, pbpX, pbpF, pbpB* | *tetA, tetO* | | *gyrA, gyrB, parB, parC, parE, prmA, prmC* |
| JCM 1112 | Human | | *ponA, pbpX, pbpF, pbpB* | *tetA, tetO* | | *gyrA, gyrB, parB, parC, parE, prmA, prmC* |
| CF48_3A | Human | | *ponA, pbpX, pbpF, pbpB* | *tetA, tetO, tetW* | | *gyrA, gyrB, parB, parC, parE, prmA, prmC* |
| MM2_3 | Human | | *ponA, pbpX, pbpF, pbpB* | *tetA, tetO, tetC* | | *gyrA, gyrB, parB, parC, parE, prmA, prmC* |
| 100_23 | Rat | | *ponA, pbpX, pbpF, pbpB* | *tetA, tetO* | | *gyrA, gyrB, parB, parC, parE, prmA, prmC* |
| I49 | Mouse | | *ponA, pbpX, pbpF, pbpB* | *tetA, tetO* | | *gyrA, gyrB, parB, parC, parE, prmA, prmC* |
| mlc3 | Mouse | | *ponA, pbpX, pbpF, pbpB* | *tetA, tetO* | | *gyrA, gyrB, parB, parC, parE, prmA, prmC* |
| lpuph | Mouse | | *ponA, pbpX, pbpF, pbpB* | *tetA, tetO* | | *gyrA, gyrB, parB, parC, parE, prmA, prmC* |
| ATCC53608 | Pig | | *ponA, pbpX, pbpF, pbpB* | *tetA, tetO* | *vanH* | *gyrA, gyrB, parB, parC, parE, prmA, prmC* |
| I5007 | Pig | | *ponA, pbpX, pbpF, pbpB* | *tetA, tetO, tetM, tetW* | | *gyrA, gyrB, parB, parC, parE, prmA, prmC* |
| ZLR003 | Pig | | *ponA, pbpX, pbpF, pbpB* | *tetA, tetO, tetW, tetL* | | *gyrA, gyrB, parB, parC, parE, prmA, prmC* |
| 20_02 | Pig | | *ponA, pbpX, pbpF, pbpB* | *tetA, tetO, tetM, tetW* | | *gyrA, gyrB, parB, parC, parE, prmA, prmC* |
| CRL1098 | Sourdough | | *ponA, pbpX, pbpF, pbpB* | *tetA, tetO* | | *gyrA, gyrB, parB, parC, parE, prmA, prmC* |
| TMW1.656 | Sourdough | | *pbpX, pbpF, pbpB, pbpA, pbpG* | *tetA, tetO* | | *gyrA, gyrB, parB, parC, parE, prmA, prmC* |
| TMW1.112 | Sourdough | | *pbpX, pbpF, pbpB, pbpA* | *tetA, tetO* | | *gyrA, gyrB, parB, parC, parE, prmA, prmC* |
| LTH5448 | Sourdough | | *ponA, pbpX, pbpF, pbpB* | *tetA, tetO* | | *parB, parC, parE, prmA, prmC* |
| LR1 | Goat | | *ponA, pbpX, pbpF, pbpB* | *tetA, tetO* | | *gyrA, gyrB, parB, parC, parE, prmA, prmC* |
| LR2 | Goat | | *ponA, pbpX, pbpF, pbpB* | *tetA, tetO* | | *gyrA, gyrB, parB, parC, parE, prmA, prmC* |
| LR3 | Goat | | *ponA, pbpX, pbpF, pbpB* | *tetA, tetO* | | *gyrA, gyrB, parB, parC, parE, prmA, prmC* |
| LR11 | Goat | | *ponA, pbpX, pbpF, pbpB* | *tetA, tetO* | | *gyrA, gyrB, parB, parC, parE, prmA, prmC* |
| LR14 | Goat | | *ponA, pbpX, pbpF, pbpB* | *tetA, tetO* | | *gyrA, gyrB, parB, parC, parE, prmA, prmC* |
| LR6 | Sheep | | *ponA, pbpX, pbpF, pbpB* | *tetA, tetO* | | *gyrA, gyrB, parB, parC, parE, prmA, prmC* |
| LR7 | Sheep | | *ponA, pbpX, pbpF, pbpB* | *tetA, tetO* | | *gyrA, gyrB, parB, parC, parE, prmA, prmC* |
| LR8 | Sheep | | *ponA, pbpX, pbpF, pbpB* | *tetA, tetO, tetM* | | *gyrA, gyrB, parB, parC, parE, prmA, prmC* |
| LR9 | Sheep | | *ponA, pbpX, pbpF, pbpB* | *tetA, tetO, tetM* | | *gyrA, gyrB, parB, parC, parE, prmA, prmC* |
| LR4 | Cow | | *ponA, pbpX, pbpF, pbpB* | *tetA, tetO* | | *gyrA, gyrB, parB, parC, parE, prmA, prmC* |
| LR10 | Cow | | *ponA, pbpX, pbpF, pbpB* | *tetA, tetO* | | *gyrA, gyrB, parB, parC, parE, prmA, prmC* |
| LR12 | Cow | | *ponA, pbpX, pbpF, pbpB* | *tetA, tetO* | | *gyrA, gyrB, parB, parC, parE, prmA, prmC* |
| LR13 | Cow | | *ponA, pbpX, pbpF, pbpB* | *tetA, tetO* | | *gyrA, gyrB, parB, parC, parE, prmA, prmC* |
| LR17 | Horse | | *ponA, pbpX, pbpF, pbpB* | *tetA, tetO* | | *gyrA, gyrB, parB, parC, parE, prmA, prmC* |
| LR18 | Horse | | *ponA, pbpX, pbpF, pbpB* | *tetA, tetO* | | *gyrA, gyrB, parB, parC, parE, prmA, prmC* |
| LR19 | Horse | | *ponA, pbpX, pbpF, pbpB* | *tetA, tetO, tetM* | | *gyrA, gyrB, parB, parC, parE, prmA, prmC* |

CFX, cefotaxime; ERM, erythromycin; PEN, penicillin; TET, tetracycline; VAN; vancomycin; CIP, ciprofloxacin. MIC, minimum inhibitory concentration.
