## Supplementary Table S3 for "Isolation and comparative genomic analysis of reuterin-producing *Lactobacillus reuteri* from poultry gastrointestinal tract"

**Supplementary Table S3.** Unique genes in the *L. reuteri* poultry human lineage VI illustrated in Figure 3.

| **Unique genes Human-Poultry VI clade** |
|---|
| Transposase IS116/IS110/IS902 family protein |
| phosphoglycerate mutase |
| Tyrosine-protein kinase YwqD |
| Capsular polysaccharide type 8 biosynthesis protein cap8A |
| ASCH domain protein |
| Transcriptional activatory protein AadR |
| Copper chaperone CopZ |
| Zinc-transporting ATPase |
| Bacterial low temperature requirement A protein (LtrA) |
| ABC transporter ATP-binding protein YxdL |
| Bacitracin export permease protein BceB |
| HTH-type transcriptional regulator ImmR |
| Phage integrase family protein |
| Putative HTH-type transcriptional regulator YwnA |
| Folate transporter FolT |
| Quinone oxidoreductase 2 |
| L-2-hydroxyisocaproate dehydrogenase |
| 2-hydroxyhexa-2-4-dienoate hydratase |
| Alpha/beta hydrolase family protein |
| Branched-chain amino acid transport protein (AzlD) |
| AzlC protein |
| Deoxyribonucleoside regulator |
| Adenosine monophosphate-protein transferase SoFic |
| GTP pyrophosphokinase YwaC |
| Response regulator MprA |
| Signal transduction histidine-protein kinase BaeS |
| Carbamoyl-phosphate synthase large chain |
| Carbamoyl-phosphate synthase small chain |
| Riboflavin biosynthesis protein RibBA |
| HTH-type transcriptional activator RhaS |
| Threonine synthase |
| Homoserine dehydrogenase |
| Homoserine kinase |
| Phosphoenolpyruvate synthase |
| Potassium lithium and rubidium/H(+) antiporter |
| High-affinity gluconate transporter |
| Glycerol-3-phosphate acyltransferase |
| Flavodoxin |
| Helix-turn-helix protein |
